## Supplemental Figures for "Parent-of-origin effects propagate through networks to shape metabolic traits"

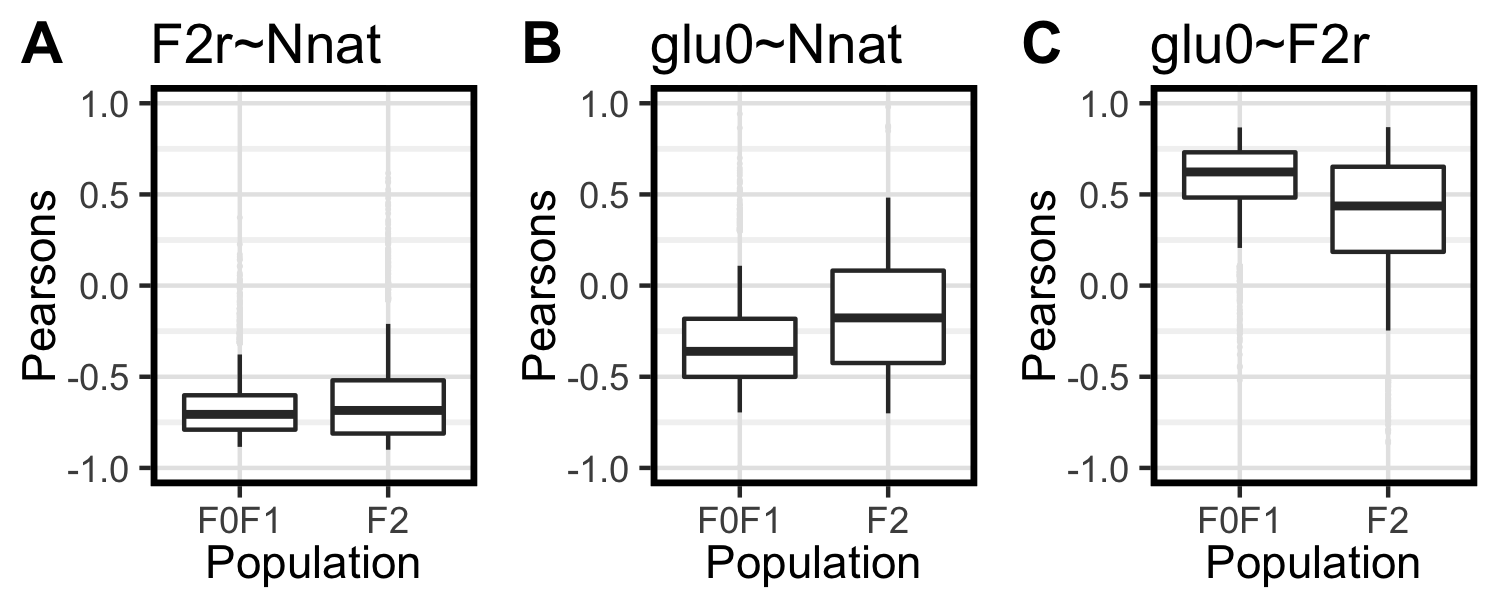

**Supplemental Figure 1: Pearson’s correlation coefficient confidence intervals.** To determine whether the pattern of correlations is significantly different between F_0_/F_1_ and F_2_ populations bootstrapping was employed (N=10, 5000 iterations). Pickets correspond to 90% confidence intervals, boxes correspond to 50% confidence intervals. Overlap of confidence intervals indicates no significant difference between populations. Correlation between F2r and Nnat expression is not significantly different (A). Correlation between Nnat and basal glucose is not significantly different (B). Correlation between F2r and basal glucose is not significantly different (C).

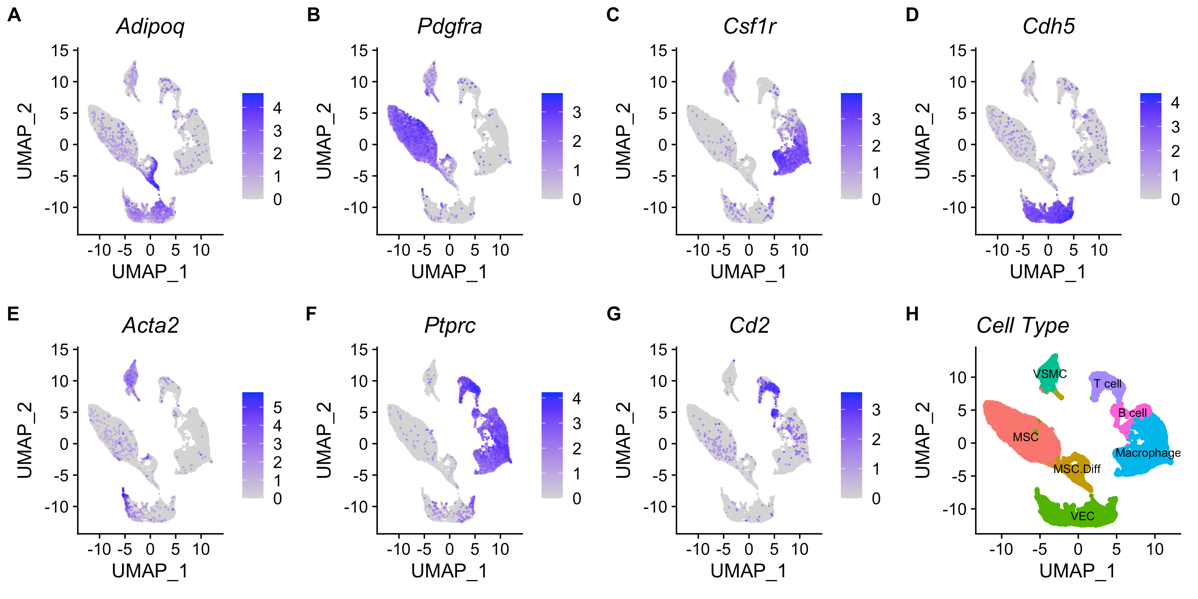

**Supplemental Figure 2. Cell types were defined using canonical markers.** Expression of these markers is shown by coloration on UMAP plots for *Adipoq* = differentiating mesenchymal stem cells (A); *Pdgfra* = mesenchymal stem cells (B); *Csf1r* = macrophage(C); *Cdh5* = vascular endothelial cells(D); *Acta2* = vascular smooth muscle cells(E); Ptprc – T cells(F); *Cd2* – B cells (G). Cell type identities were assigned by expression of these genes (H)

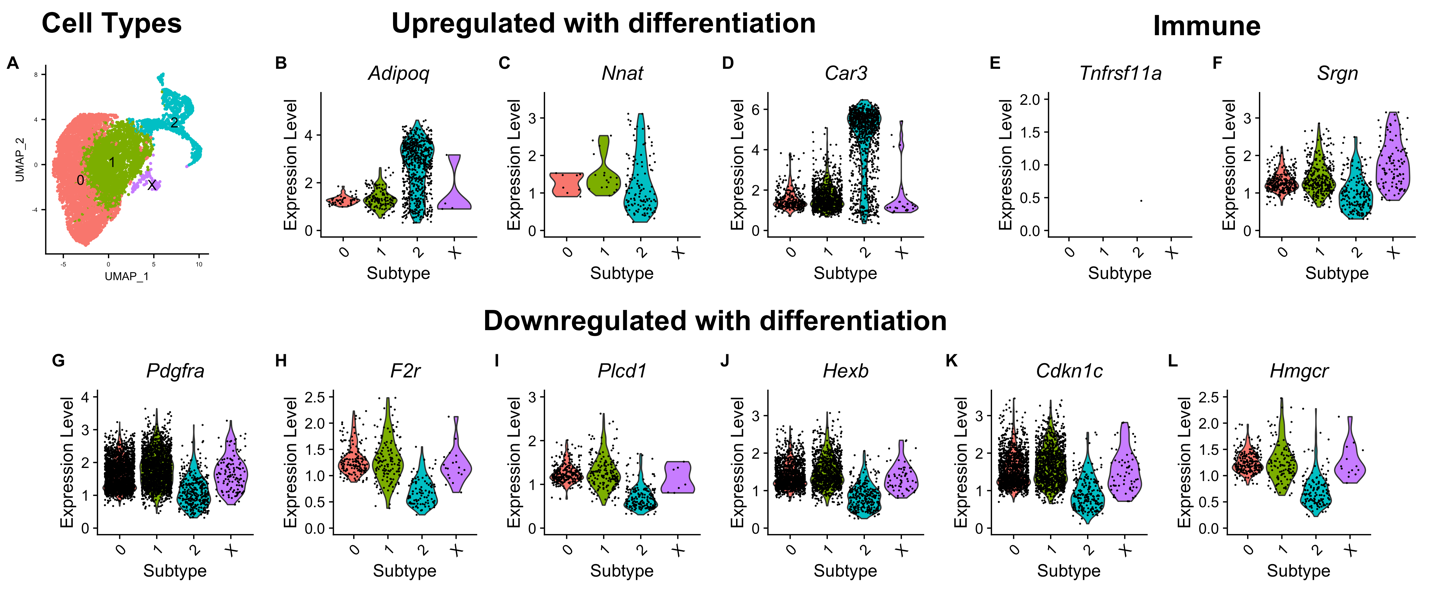

**Supplemental Figure 3. Candidate genes are differentially expressed across adipogenic trajectory.** Cells along the adipogenic trajectory were clustered into 4 groups [0, 1, 2, X](A). Adipogenic trajectory was defined by the expression of Adipoq in differentiation MSC (B) and by Pdgfra in undifferentiated MSC (G). Expression of candidate genes identified by integrating F0/F1 RNAseq with F16 data is shown. Only cells with non-zero expression are plotted. Two candidate genes are primarily expressed by immune cells (E & F), but one of which is differentially expressed along the adipogenic trajectory (F). Two candidate genes are primarily expressed along the trajectory and are upregulated with differentiation (C & D). Five candidate genes are primarily expressed along the trajectory and downregulated with differentiation (H-L).

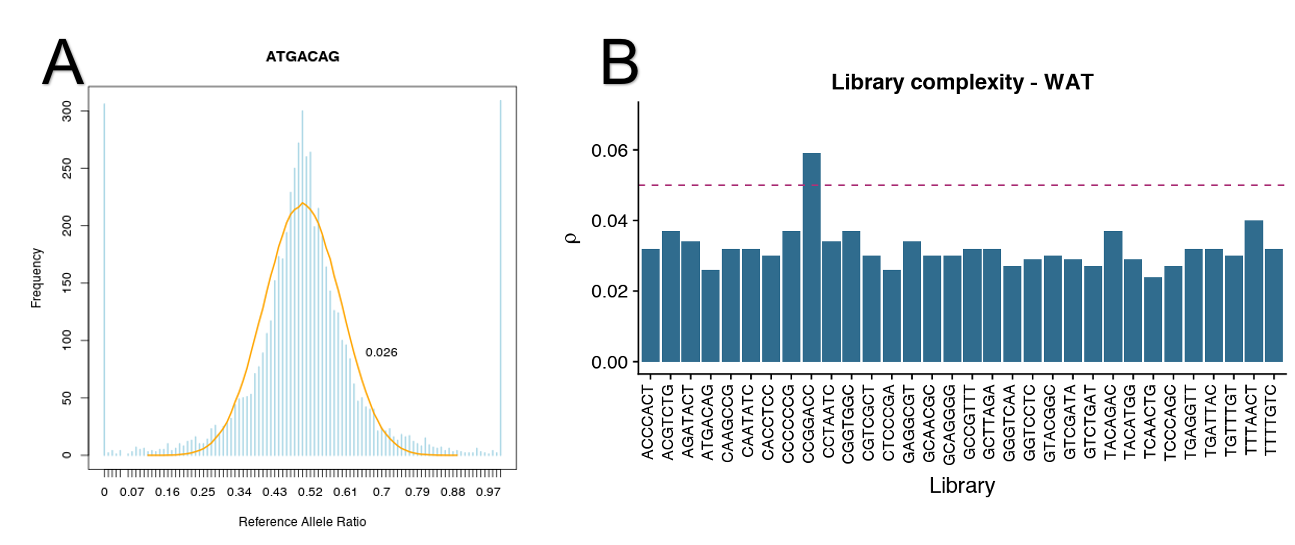

**Supplemental Figure 4: RNAseq libraries are sufficiently complex to detect allele specific expression.** Complexity was measured by fitting a beta-binomial distribution to the distribution of L_bias_ values. A representative distribution (**A**) of L_bias_ values (blue) and fit beta-binomial (orange). The shape parameters (α, β) of the beta-binomial distribution are used to calculate dispersion (ρ). Dispersion value less than 0.05 indicate sufficient complexity. One library was not sufficiently complex (**B**) and was removed from analyses.

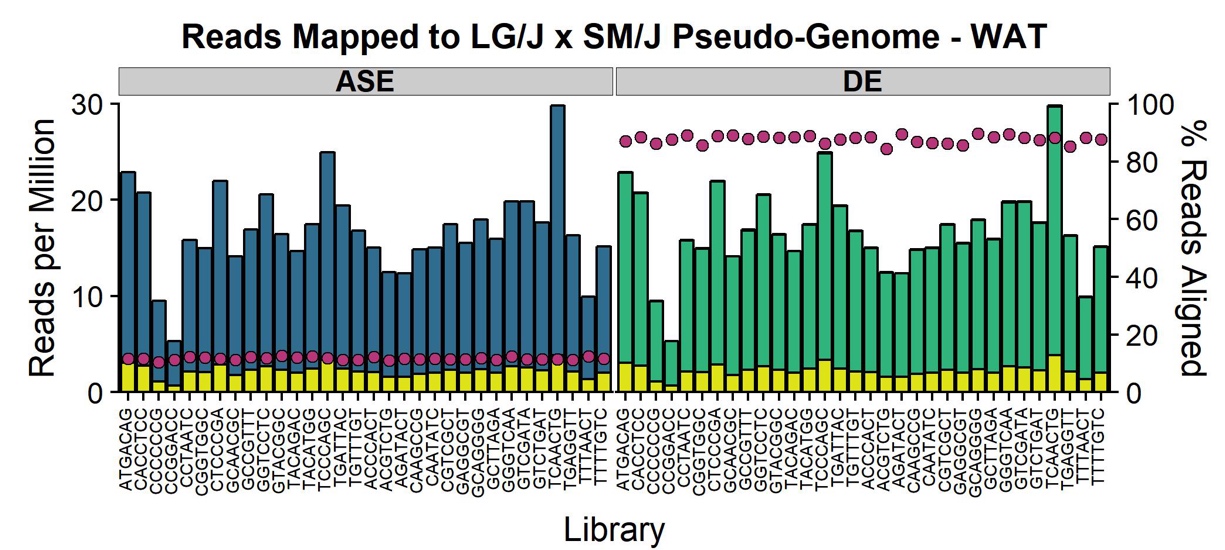

**Supplemental Figure 5: Number of reads mapped to LG/J x SM/J pseudo-genome.** We isolated total RNA from white adipose (n = 32 per sex/diet/cross cohort) and constructed RNA-Seq libraries. Libraries were sequenced 1 sample per lane on an Illumina HiSeq 4000 (Illumina, San Diego, USA) using 2x100bp PE reads. FASTQ files were filtered to remove low quality reads. Remaining reads were aligned against both LG/J and SM/J pseudo-genomes simultaneously using STAR with either multimapping disallowed (allele-specific expression, **ASE**) or multimapping allowed (differential expression, **DE**). The stacked bars denotes the number of reads per million (left y-axis) and are color-coded by category (uniquely = yellow, multi-loci = green, too many = blue). For the ASE alignment, “too many” reads are defined as n > 1 since multimapping was disallowed. For the DE alignment, “multi-loci” reads are defined as 1 < n < 10 and “too many” reads are defined as n > 10. The pink dots denote the primary alignment rates (right y-axis) for each library. Alignment rates are defined as (# reads mapped uniquely + # reads mapped multi-loci / total # input reads).

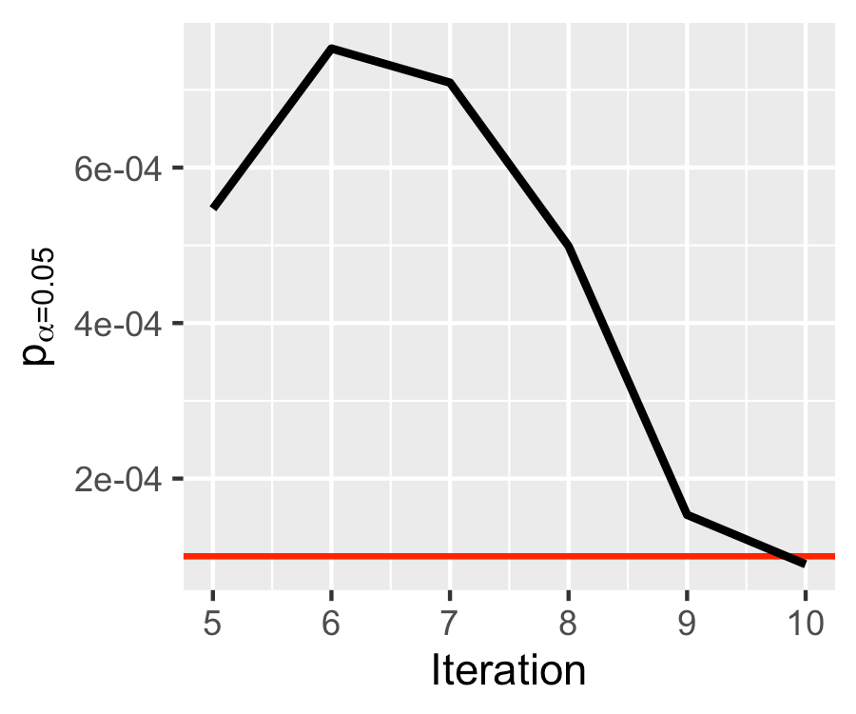

**Supplemental Figure 6: Stable null permutation plots for allele-specific expression.** Context was permuted to generate an appropriate null model of p-values. Each iteration represents one genome wide permutation where every gene (7,869) was permuted separately, thus each iteration represents 7,869 separate shuffles. Null distribution stability was evaluated by calculating the critical value (in this case a significance estimate) for alpha = 0.05 at each iteration. The standard deviation of critical values was calculated after each iteration for the last 5 iterations. The model was considered stable once variation in critical values dropped below 1e-4 (red line). The variation in critical values of the null model approaches zero over ten genome wide iterations. This approximation towards zero demonstrates the stabilization of the null model. By ten genome wide shuffles the critical value fluctuates <1e-4 units per iteration in the most complex model, meaning that at false discovery rate estimates might vary by ± 1e-4.

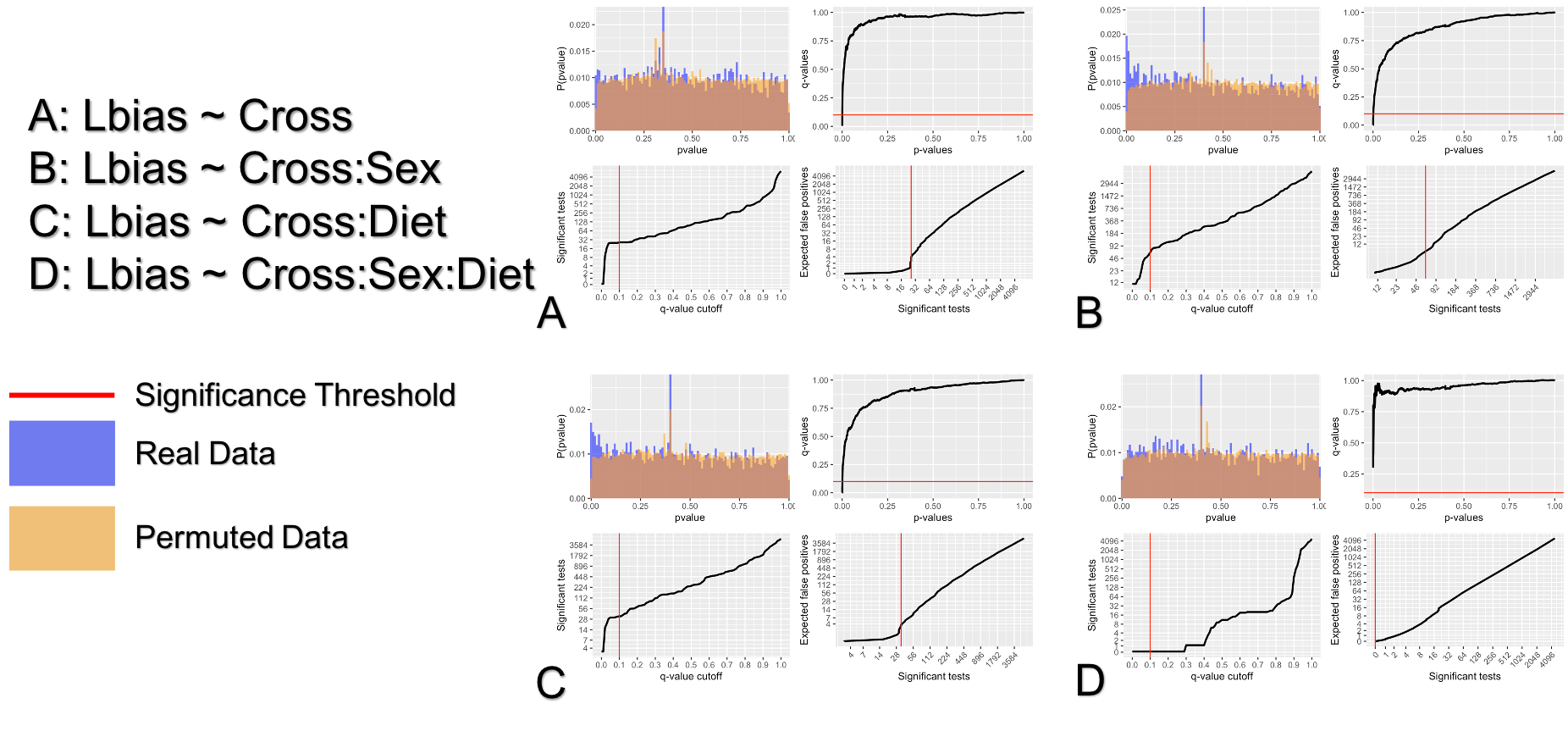

**Supplemental Figure 7: Multiple tests correction of ASE detection.** Multiple tests correction was performed by estimating the false discovery rate using a permuted null model of p-values. Context was permuted to generate an appropriate null model of p-values. The distribution of permuted (orange) and real (blue) p-values is shown for each term in the model in the top left of each subplot. False discovery rate was estimated as the proportion of tests below a given significant threshold under the null relative to the real data (Null/Real). A ratio of 1 meaning that ~100% of observations at that significance threshold are likely false discoveries. The relationship between false discovery rate and p-value threshold is shown in the top right subplot for each mode. The number of significant tests at a given acceptable false discovery rate is shown as the bottom left subplot for each model. The number of false positives relative to the number of significant tests is shown as the bottom right subplot for each model.

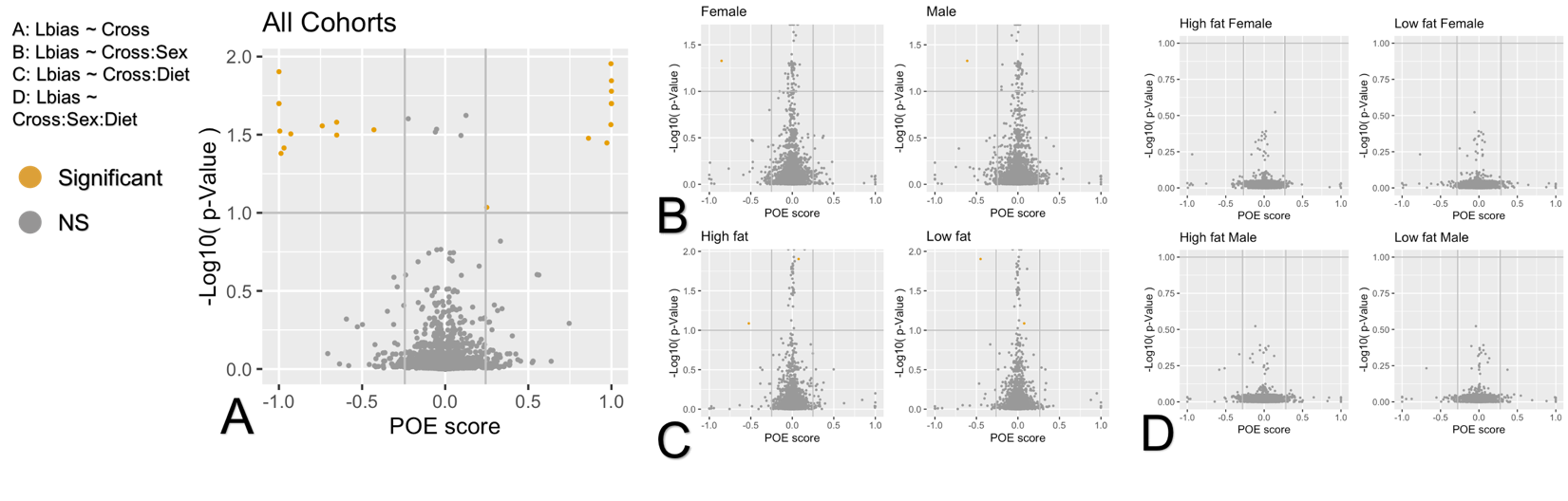

**Supplemental Figure 8: Volcano plots of parent-of-origin dependent allele specific expression.** Log_10_ transformed FDR plotted against POE score [ mean SxL Lbias – mean LxS Lbias ] for each context. Genes passing both POE score and significance thresholds are shown in orange. Genes failing to meet these criteria are shown in grey.

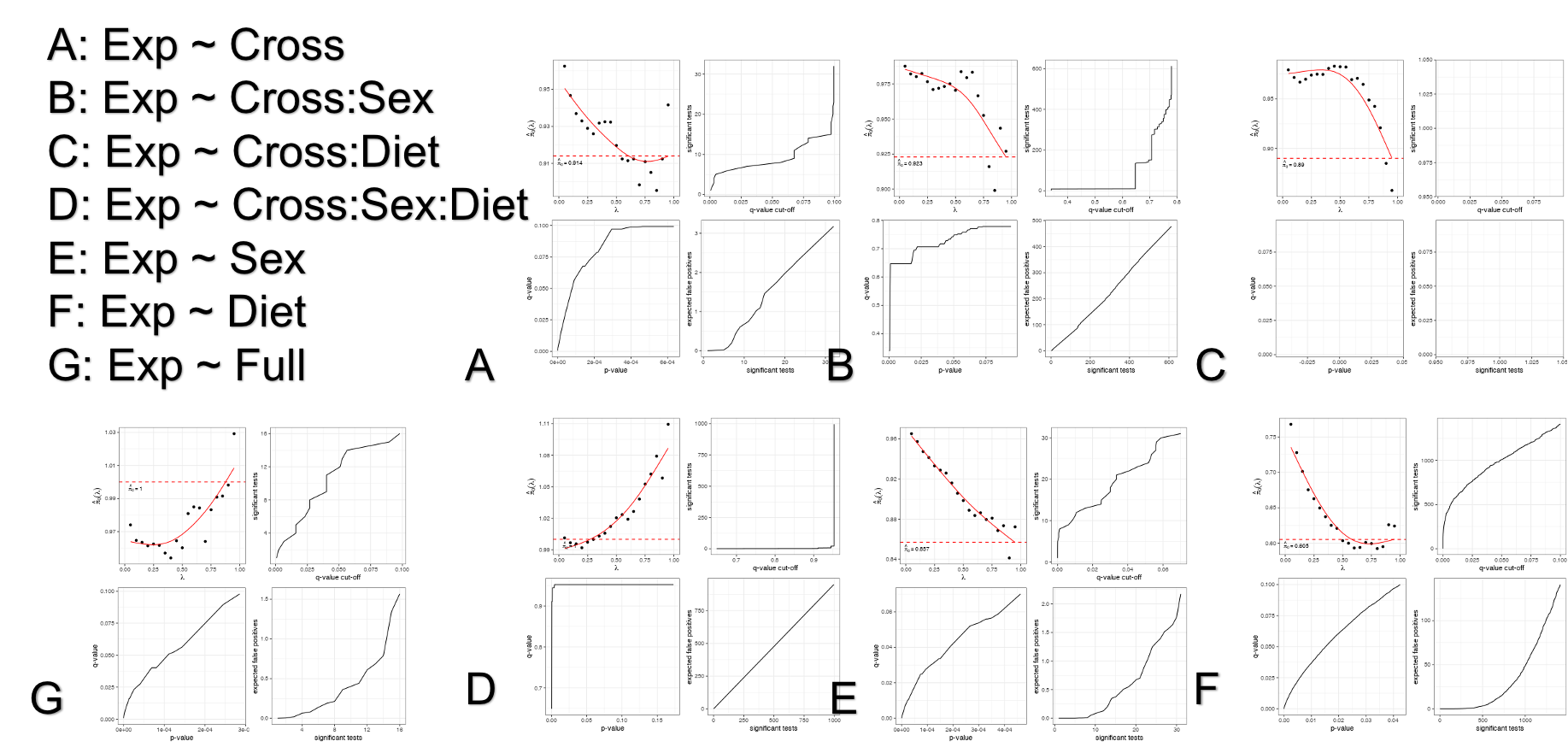

**Supplemental Figure 9: Multiple tests correction of DE detection.** Multiple tests correction was performed by estimating the false discovery rate using the “qvalue” R package without the generation of a permuted null model. The underlying null model is selected by the model by controlling $\hat{\pi_{0}}$. False discovery rate (q-value) is estimated as the proportion of tests below a given significant threshold derived from the estimated null model relative to the real model (Null/Real). A ratio of 1 meaning that ~100% of observations at that significance threshold are likely false discoveries. The selection of $\hat{\pi_{0}}$ is shown in the top left of each subplot for each model. The relationship between false discovery rate and p-value threshold is shown in the bottom left subplot for each mode. The number of significant tests at a given acceptable false discovery rate is shown as the top right subplot for each model. The number of false positives relative to the number of significant tests is shown as the bottom right subplot for each model.

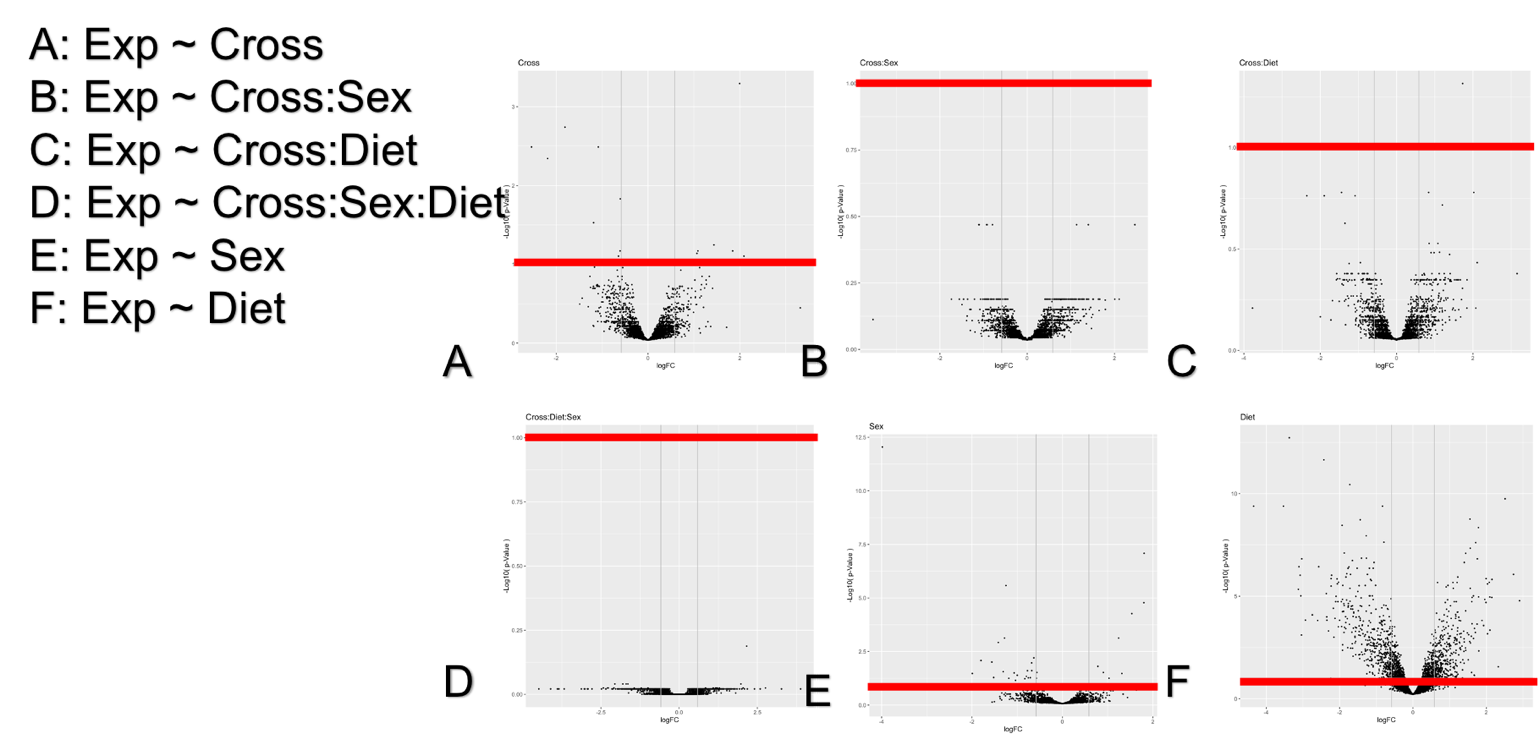

**Supplemental Figure 10: Volcano plots of differentially expressed genes.** Log_10_ transformed FDR plotted against log_2_ transformed fold change for each context. The FDR threshold we used was 0.1 (red line).

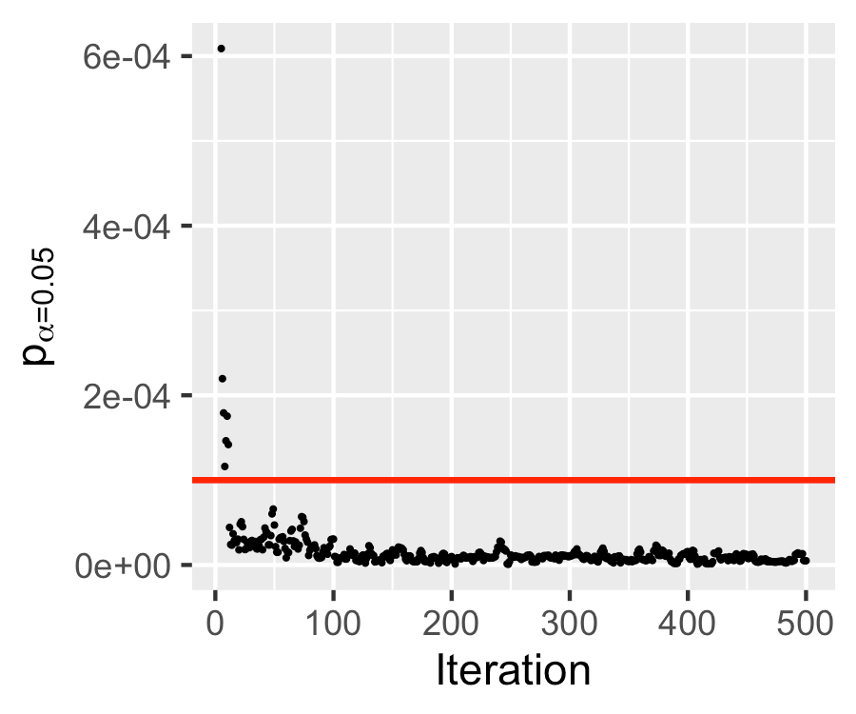

**Supplemental Figure 11: Stable null permutation plots for network pairs**. Context and expression were permuted to generate an appropriate null distribution of p-values. Significance for each pair was estimate by likelihood ratio test using a chi-squared approximation. Null distribution stability was evaluated by calculating the critical value (in this case a significance estimate) for alpha = 0.05 at each iteration. The standard deviation of critical values was calculated after each iteration for the last 5 iterations. The model was considered stable once variation in critical values dropped below 1e-4 (red line). The variation in critical values under the null model approaches zero over 100 iterations. By 500 shuffles the critical value fluctuates well below 1e-4 units per iteration in the most complex model, meaning that at worst FDR estimates would vary by well below ± 1e-4.

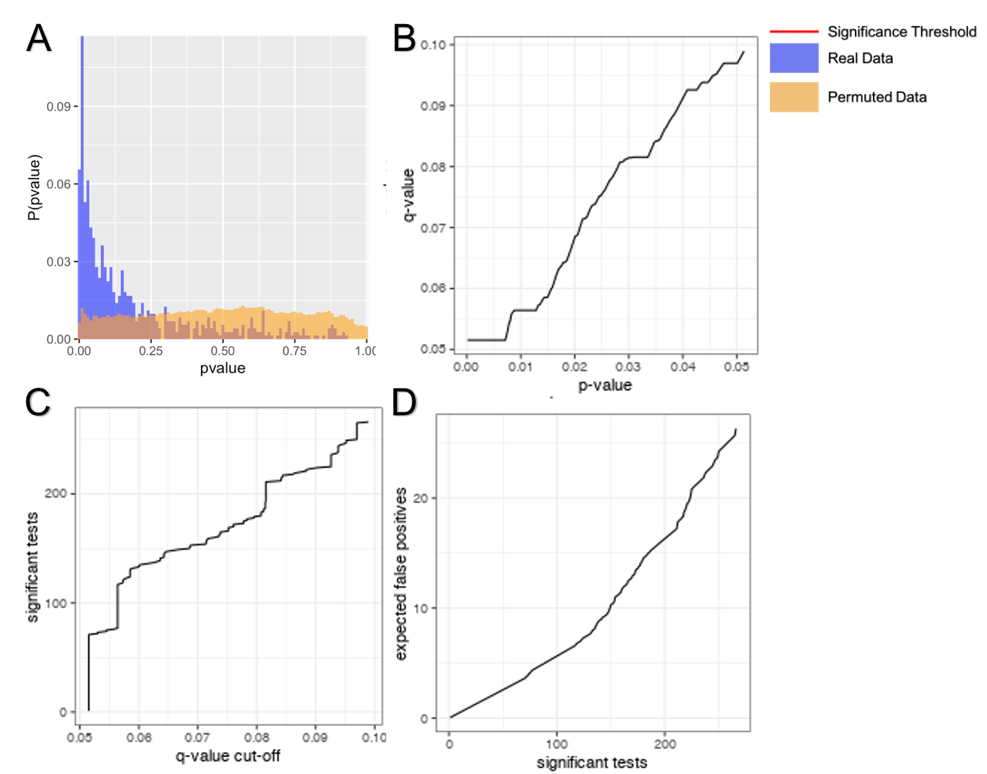

**Supplemental Figure 12: Multiple tests correction of pairwise network construction.** Multiple tests correction was performed by estimating the false discovery rate using a permuted null model of p-values. The distribution of permuted (orange) and real (blue) p-values is shown (A). False discovery rate was estimated as the proportion of tests below a given significant threshold under the null relative to the real data (Null/Real). A ratio of 1 meaning that ~100% of observations at that significance threshold are likely false discoveries. The relationship between false discovery rate and p-value threshold is shown (B). The number of significant tests at a given acceptable false discovery rate is shown (C). The number of false positives relative to the number of significant tests is shown (D).

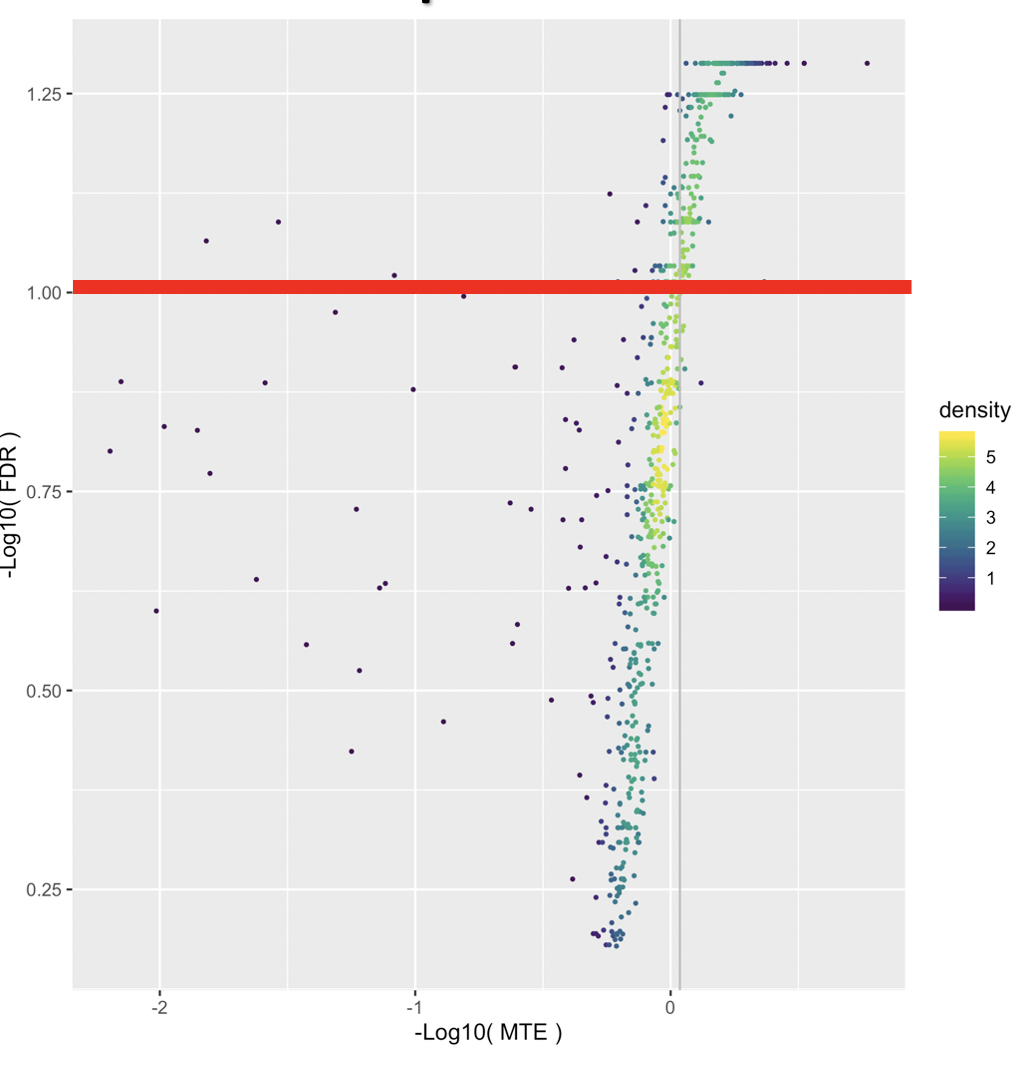

**Supplemental Figure 13: Volcano plots of network construction.** Log_10_ transformed FDR is plotted against log_10_ transformed mean test error (MTE). The FDR threshold value of 0.1 is shown as the red line. The MTE threshold was set to the upper 1th percentile of the permuted model MTE and is show as the grey vertical line. Each gene is represented by a single point. Coloration of points is determined by the joint FDR/MTE density.

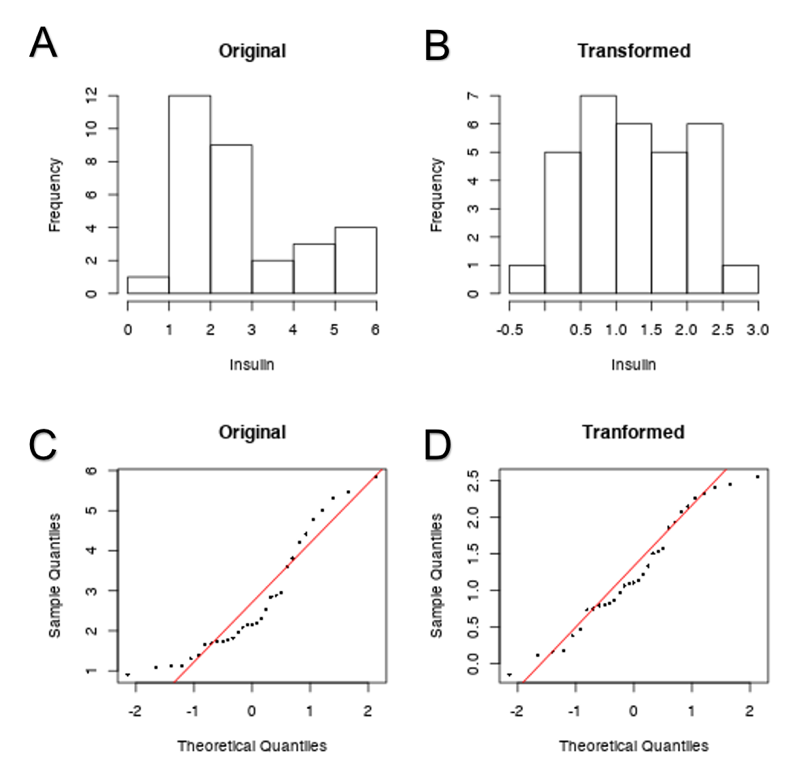

**Supplemental Figure 14: Example transformation of F_1_ phenotypes.** In order to meet the assumptions of normality required by Pearson’s correlation test, non-normally distributed phenotypes were log transformed. Normality was evaluated by Shapiro-Wilkes test (p<0.05 indicates non-normality). Serum insulin is one such phenotype. The probability distribution of serum insulin is shown before (A) and after transformation (B). This distribution is also compared to a normal distribution via quantile-quantile plot before (C) and after (D) transformation.

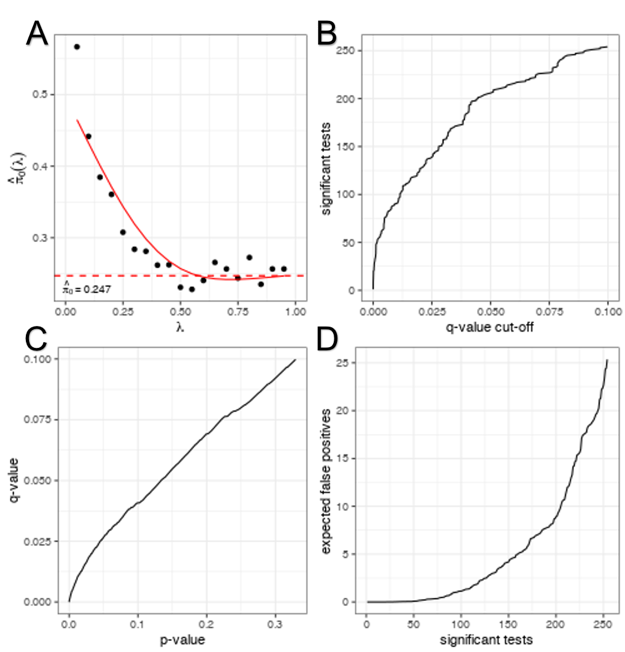

**Supplemental Figure 15: Multiple tests correction of phenotype correlations.** Multiple tests correction was performed by estimating the false discovery rate using the “qvalue” R package without generating a permuted null model. The underlying null model is selected by controlling $\hat{\pi_{0}}$. False discovery rate (q-value) is estimated as the proportion of tests below a given significant threshold derived from the estimated null model relative to the real model (Null/Real). A ratio of 1 meaning that ~100% of observations at that significance threshold are likely false discoveries. Selecting $\hat{\pi_{0}}$ (A) allows us to determine the relationship between false discovery rate and p-value threshold (C). Using this relationship, we can determine the number of significant tests at a given acceptable false discovery rate (B). From this we can determine the number of false positives relative to the number of significant tests (D).

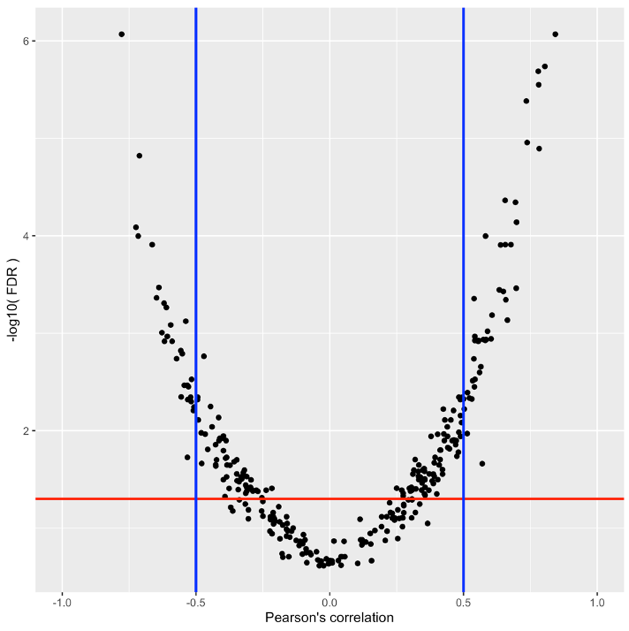

**Supplemental Figure 16: Volcano plots of phenotypes correlated with POE net gene expression.** Log_10_ transformed FDR plotted against Pearson’s correlation coefficient. Each dot represents a phenotype/gene pair. The FDR threshold was set to 0.1 (red line). The correlation coefficient threshold was set to |0.5| (blue line).

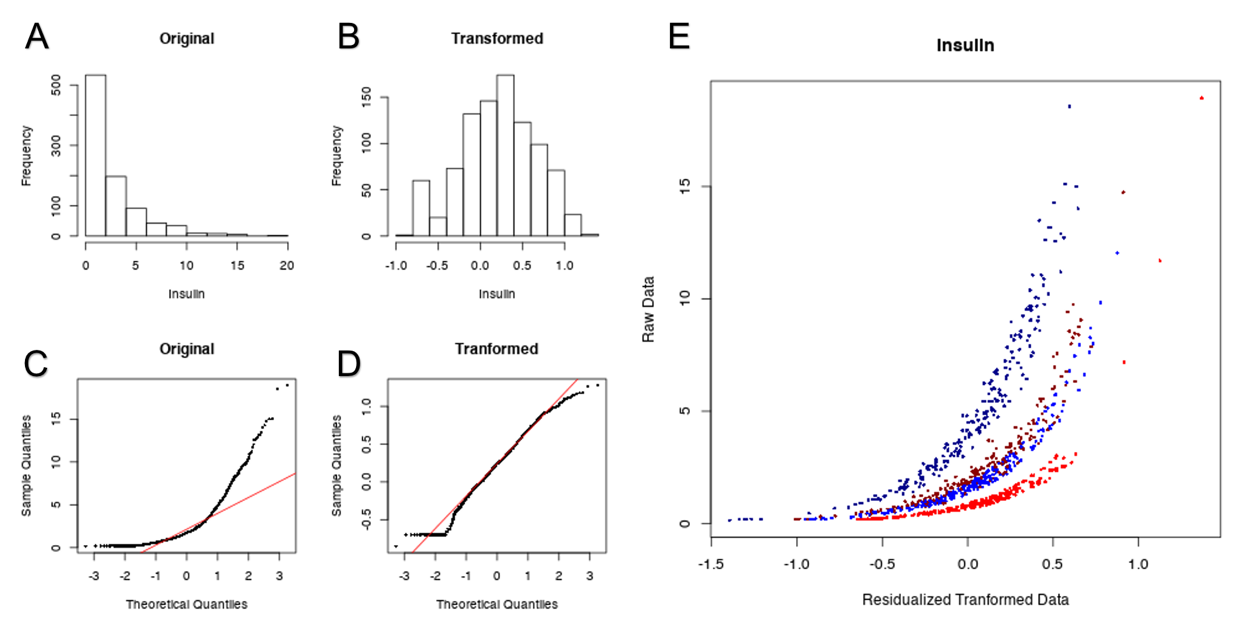

**Supplemental Figure 17: Example transformation of F_16_ phenotypes.** In order to meet the assumptions of normality required by ANOVAs, non-normally distributed phenotypes were log transformed and residualized to remove the effects of sex and diet. Normality was evaluated by Shapiro-Wilkes test (p<0.05 indicates non-normality). Serum insulin is one such phenotype. The probability distribution of serum insulin is shown before (A) and after transformation (B). This distribution is also compared to a normal distribution via quantile-quantile plot before (C) and after (D) transformation. The removal of sex and diet effects on serum insulin are shown (E) for high fat males (dark blue), low fat males (light blue), high fat females (dark red), low fat fed females (light red).

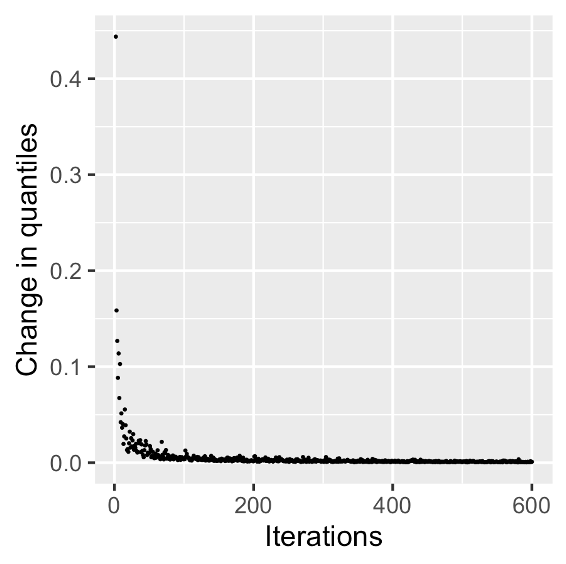

**Supplemental Figure 18: Stable null permutations plot for epistasis**. Imprinting scores were permuted to generate an appropriate null model p-value. Null distribution stability was evaluated by calculating the total quantile deviation (TQD) at each iteration. The TQD of F-statistics under the null model approaches zero over 600 iterations.

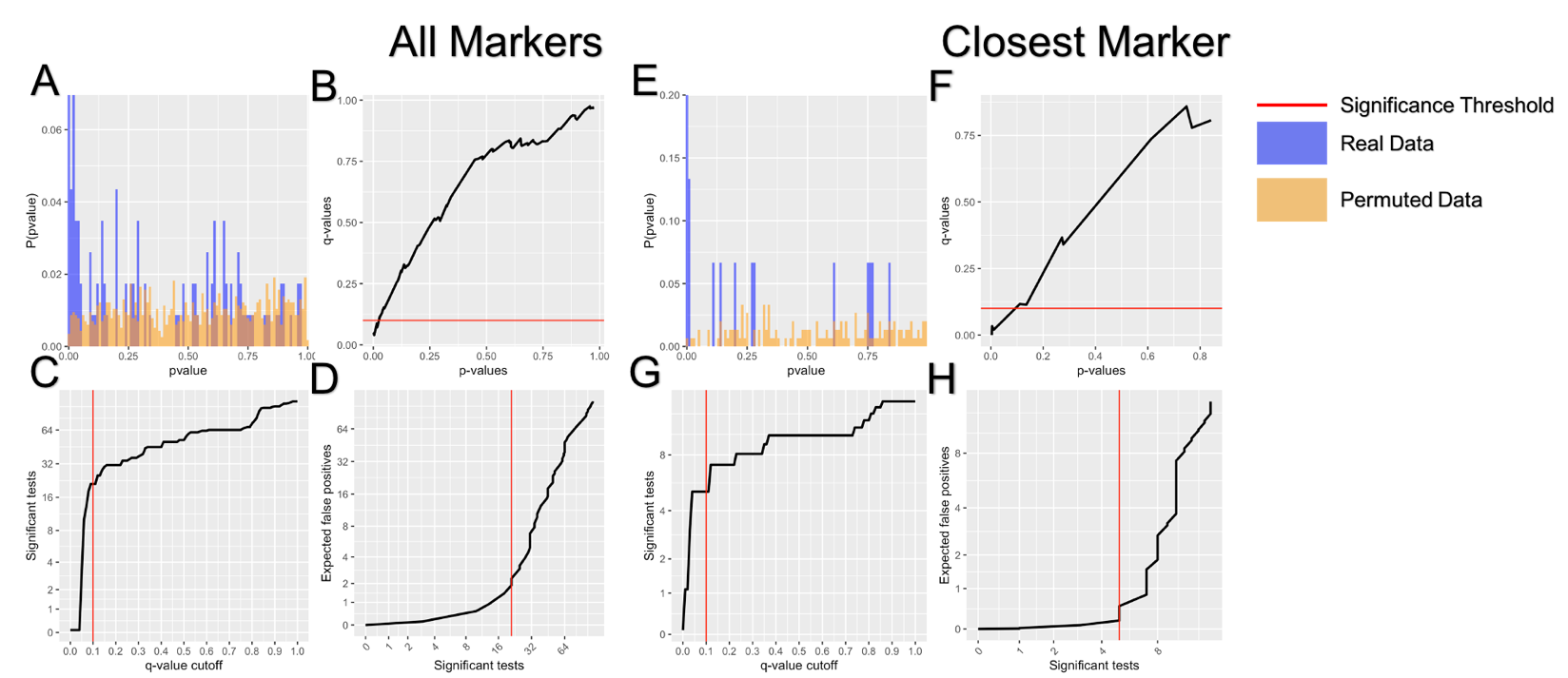

**Supplemental Figure 19: Representative multiple tests correction of imprinting:imprinting epistasis.** Multiple test corrections was performed on each phenotype individually. Here were show this for the GTT area under the curve at 10 weeks phenotype using both all markers within 1.5Mb (A-D) and only the nearest marker (E-H). Multiple tests correction was performed by estimating the false discovery rate using a permuted null model of p-values. Imprinting scores were permuted to generate an appropriate null model p-values. The distribution of permuted (orange) and real (blue) p-values is shown (A & E). False discovery rate was estimated as the proportion of tests below a given significant threshold under the null relative to the real data (Null/Real). A ratio of 1 meaning that ~100% of observations at that significance threshold are likely false discoveries. The relationship between false discovery rate and p-value threshold is shown (B & F). The number of significant tests at a given acceptable false discovery rate is shown (C & G). The number of false positives relative to the number of significant tests is shown (D & H).

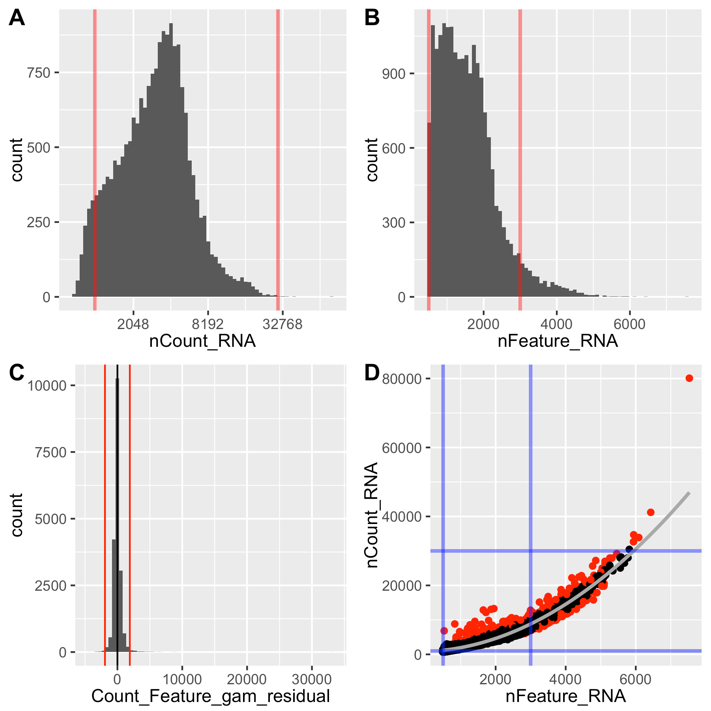

**Supplemental Figure 20. Single cell quality was controlled.** The number of counts for a given gene in a given cell nCount (A), the number of genes expressed in a given cell nFeature (B), and their covariation (C) is predictive of cell quality. The strength of their relationship was measured as the residual for each cell in a generalized additive model predicting nCount from nFeature. Thresholds for these metrics (red/blue lines) were used to decide which cells to include. Cells with unacceptably high residuals (red dot) were thrown out.

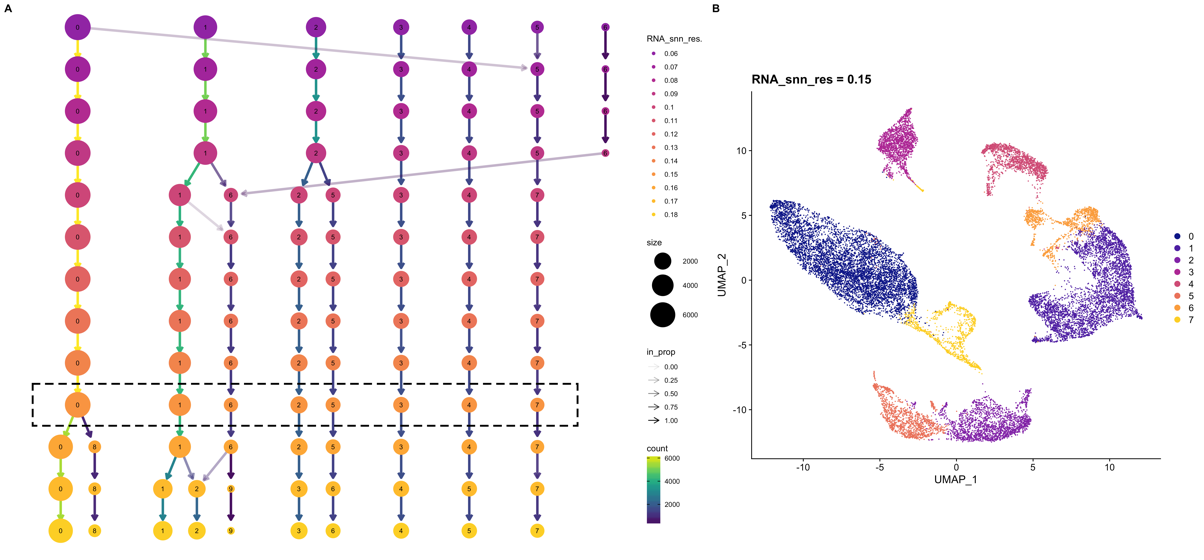

**Supplemental Figure 21. Determining the resolution for clustering.** Dimensionality reduction of high dimensional single-cell RNAseq data requires the selection of a resolution parameter. This parameter is selected by seeing how stable cluster number and assignment is across a range of resolution parameter values as shown on a cluster tree(A). Here we selected the highest resolution parameter that stably clustered the expected cell populations. The selected value was 0.15. Using this value, dimensionality reduction is visualized using the UMAP method (B). Cluster identifiers is consistent between cluster tree and UMAP projection.

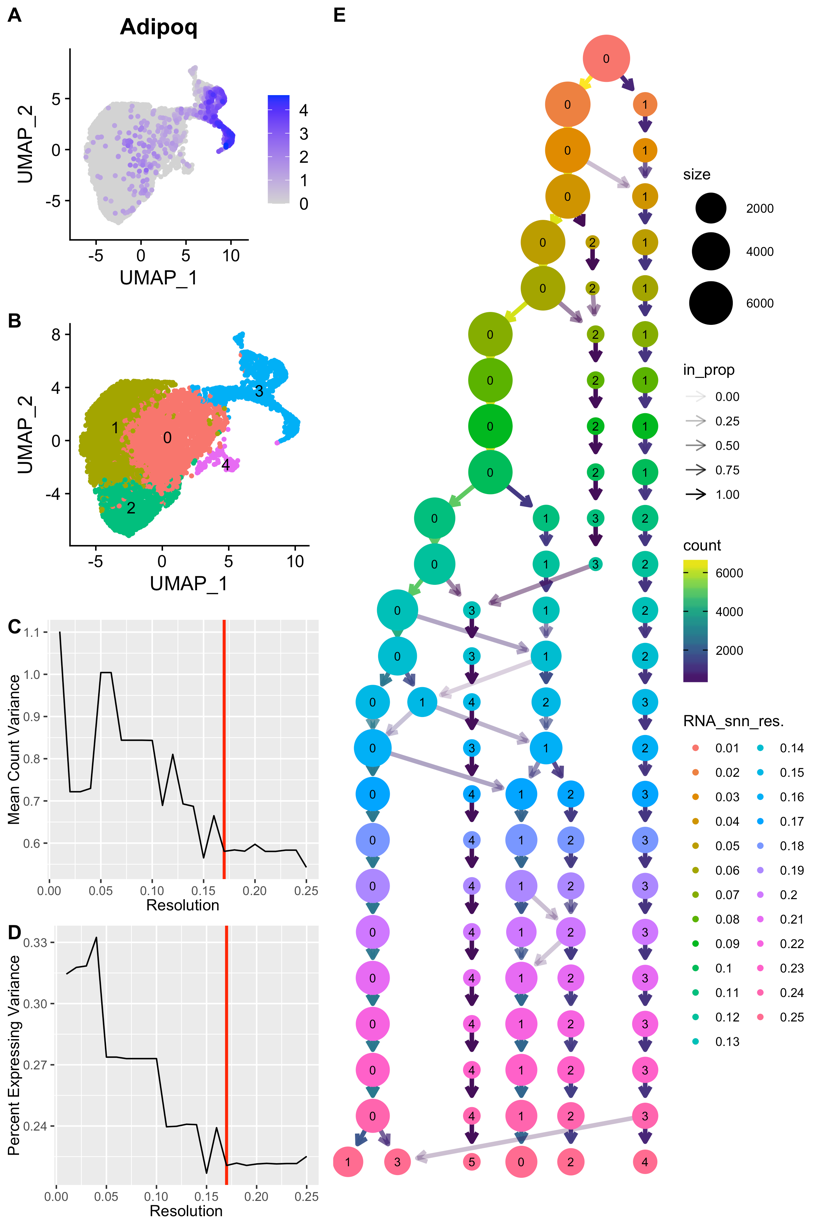

**Supplemental Figure 22. Adipocyte clustering resolution was selected to minimize Adipoq variation.** We sought to test for differential expression of genes across the adipogenic trajectory. Cells from the trajectory were subset and re-clustered. We sought to cluster cells into groups that best corresponded to the different stages of differentiation (B). To this end we used Adipoq expression as a marker of differentiation (A). This dimensionality reduction of high dimensional single-cell RNAseq data requires the selection of a resolution parameter. We tested a range of resolution parameters from 0.01 to 0.25 and observed how resolution affected clustering (E). For each resolution parameter the variation in mean variation in non-zero Adipoq expression within all clusters was calculated (C). The same was done to calculate the mean variation in percent of cells with expression >0 within all clusters. By finding the clustering conditions that in which Adipoq expression was stably minimized, we were able to select the resolution parameter that best clustered cells into the different stages of differentiation (resolution=0.17). Dimensionality reduction is visualized using the UMAP method (B). Cluster identifier is consistent between cluster tree and UMAP projection.
